# Not-so-working memory: Drift in fMRI pattern representations during maintenance predicts errors in a visual working memory task

**DOI:** 10.1101/341990

**Authors:** Phui Cheng Lim, Emily J. Ward, Timothy J. Vickery, Matthew R. Johnson

## Abstract

Working memory (WM) is critical to many aspects of cognition, but it frequently fails. Much WM research has focused on capacity limits, but even for single, simple features, the fidelity of individual representations is limited. Why is this? We used fMRI and a pattern-based index of “representational drift” to investigate how ongoing changes in brain activity patterns throughout the WM maintenance period predicted performance, using a delayed-match-to-sample task for a single item with a single critical feature: orientation. In trials where the target and probe stimuli matched, participants incorrectly reported more non-matches when their activity patterns drifted away from the target. In trials where the target and probe did not match, participants incorrectly reported more matches when their activity patterns drifted towards the probe. Our results suggest that WM errors are not simply due to unstructured noise, but also drift within representation space that can be indexed by neuroimaging.

## INTRODUCTION

Working memory (WM) is critical to many aspects of cognition and behavior, yet it is far from perfect. Significant research effort has been dedicated to investigating the limits of WM capacity, both in terms of number of items (e.g., Awh et al., 2007; Luck and Vogel, 1997; Vogel et al., 2001; Xu and Chun, 2006) and in terms of the informational complexity of those items (e.g., Alvarez and Cavanagh, 2004; Xu and Chun, 2006). However, even when the number of items and their complexity are within their nominal limits, WM failures are common. How many times a day do we walk into a room and forget what we came in for? How often do we have to search for an item that we could swear we just left on the kitchen table, only to find out it was actually sitting on the couch? Despite the ubiquity of such WM failures in everyday life, the mechanisms underlying them are relatively poorly understood. If we are not exceeding the design specifications of the human working memory system in terms of item number or complexity, why doesn’t working memory work perfectly all the time?

Although the bulk of research on WM failures has focused on exceeding capacity limitations in some form (Luck and Vogel 1997, 2013; Vogel et al. 2001; Alvarez and Cavanagh 2004; Zhang and Luck 2008; Bays et al. 2009), or on interference from external factors such as distractor items (Kim et al. 2005; Vogel et al. 2005; Yoon et al. 2006; Derrfuss et al. 2017), a number of other psychological and neural mechanisms have been proposed to underlie WM performance, even in cases where loads are low and no explicit distractors are present. In terms of neural factors, WM has been studied extensively with neuroimaging techniques, and task performance has been linked to different aspects of brain activity during WM. For instance, both electroencephalography (EEG) and functional magnetic resonance imaging (fMRI) studies have found that successful WM performance is linked to greater power and/or synchrony in certain frequency bands (Khader et al. 2010; Balsters et al. 2013; Solomon et al. 2017). Several other fMRI studies have documented above-baseline levels of brain activity during the WM maintenance period and found that the amount of activity during this period corresponds with greater recall on a subsequent long-term memory test (Brewer et al. 1998; Wagner et al. 1998; Ranganath et al. 2005; Blumenfeld and Ranganath 2006). Although a number of other fMRI studies have investigated the relationship between brain activity during WM maintenance and performance on the WM task itself, they found no or limited suprathreshold activation differences during WM maintenance between accurate and inaccurate WM responses (Hannula and Ranganath 2008; Bergmann et al. 2015, 2016). One potential explanation comes from behavioral and modeling evidence suggesting that WM failures may occur as a result of “drift” in neural population activity, wherein mental representations become less accurate over time due to the accumulated effects of neural noise (Schneegans and Bays 2018) or to representational distortions (Lupyan 2008), which do not necessarily entail a change in overall activity levels. However, this hypothesis has not previously been tested directly in humans via neuroimaging.

As noted, previous fMRI studies focusing on the aggregate activity of entire brain regions may not have been sufficiently sensitive to measure fluctuations in the quality or fidelity of the information encoded by that activity, which is a necessary prerequisite for directly testing the drift hypothesis. However, research in recent years has significantly advanced our understanding of how WM representations may be encoded in more fine-grained brain activity patterns corresponding to specific memoranda, rather than by a region’s overall activation. Several studies have reported re-instantiation of perceptual brain activity patterns during WM (or during closely related activities such as mental imagery, which activate similar representations; Albers et al., 2013), wherein activity patterns exhibited during maintenance of the remembered stimulus reflect those observed during initial perception of that stimulus (Harrison and Tong 2009; Lee et al. 2012; Albers et al. 2013; LaRocque et al. 2013; Johnson and Johnson 2014). Other research examining brain activity patterns during WM maintenance has found that task-irrelevant features are encoded less strongly, if at all, compared to task-relevant features (Serences et al. 2009; Jackson et al. 2017), suggesting that these activity patterns are not merely passive re-creations of perceptual stimuli, but rather reflect an active process wherein observers prioritize storage of relevant information. Given these findings, similar pattern-based analyses may provide a valuable tool for testing hypotheses related to drift in WM representations and its relation to WM performance.

Supporting the putative utility of this analytic approach, some previous neuroimaging research has indeed found that pattern similarity can be used to infer the fidelity of WM representations and predict how well items are later remembered. While much of the extant research relating the quality of brain activity patterns to superior memory performance has focused on the link between WM and long-term memory (Xue et al. 2010; Kuhl et al. 2011, 2012; Ward et al. 2013), there is some evidence that the fidelity of memory representations during WM maintenance affects WM performance as well (Ester et al. 2013; Sprague et al. 2014). For instance, in an individual-differences study, Ester and colleagues analyzed orientation-selective responses in visual cortex and created individual tuning profiles for each participant that were predictive of task performance, suggesting that the relative “quality” of each participant’s WM representations was indicative of that person’s memory acuity. However, that study only examined individual differences, and did not determine whether the quality of representations also predicted WM performance within subjects.

Thus, significant evidence exists to support a link between brain activity during WM maintenance and subsequent behavior, although the exact nature of the relationship between maintenance activity patterns and WM performance has not been thoroughly explored. In other words, given that brain activity patterns during WM maintenance of a particular stimulus appear to mirror those observed during visual perception of that stimulus (Harrison and Tong 2009; Albers et al. 2013; Johnson and Johnson 2014) and that pattern similarity during encoding and maintenance is linked to subsequent long-term memory performance (Xue et al. 2010; Kuhl et al. 2011; Ward et al. 2013), it seems plausible that fMRI pattern similarity could be used to test the “drift hypothesis”; namely, that fluctuations in neural activity patterns during maintenance could lead to incorrect performance during a subsequent WM probe, even with successful encoding.

The present fMRI study therefore investigates how brain activity patterns associated with specific stimuli, particularly during the maintenance period, might predict the accuracy of performance on a later WM probe. We used a variation on a classic delayed match-to-sample (DMTS) task with oriented Gabor patch stimuli that has been used in past studies of brain activity patterns during WM maintenance (Harrison and Tong 2009), but with modifications allowing us to track changes in the fidelity of neural pattern representations throughout each trial. We refer to fluctuations in pattern representations as *representational drift* and relate those changes to the probability of successful versus unsuccessful WM performance in a task with nominally low demands (one item, one critical feature) on WM capacity.

## MATERIALS AND METHODS

### Participants

Twenty self-reported healthy young adults [9 females, 18 right-handed, mean age 25.1 years ±4.2 (SD)] participated in exchange for monetary compensation. All had normal or corrected-to-normal vision and provided informed consent. Procedures were approved by the Institutional Review Board at the University of Nebraska-Lincoln. Seven additional participants also took part in the study, but their datasets were rejected because of either excessive head motion or low accuracy on the behavioral task (<60% combined accuracy across all runs).

### Delayed-Match-To-Sample (DMTS) task

#### Procedure

Participants completed seven runs of the delayed match-to-sample (DMTS) task, comprising one initial pre-scan run to calibrate task difficulty (see Staircasing) and six runs in the scanner. Each run had 24 trials, lasting 24 seconds each (9.6 minutes total per run). On each trial (see **Figure 1A**), the target stimulus (a Gabor patch; for details, see Stimuli) was first presented for 1s at the center of the display against a 50% gray background. This was immediately followed by a series of briefly presented mask images, slightly larger than the stimulus, presented for a total of 1s. The mask interval was followed by a fixation interval of 10s, during which participants maintained fixation on a small white dot at the center of the screen.

**Figure 1.**
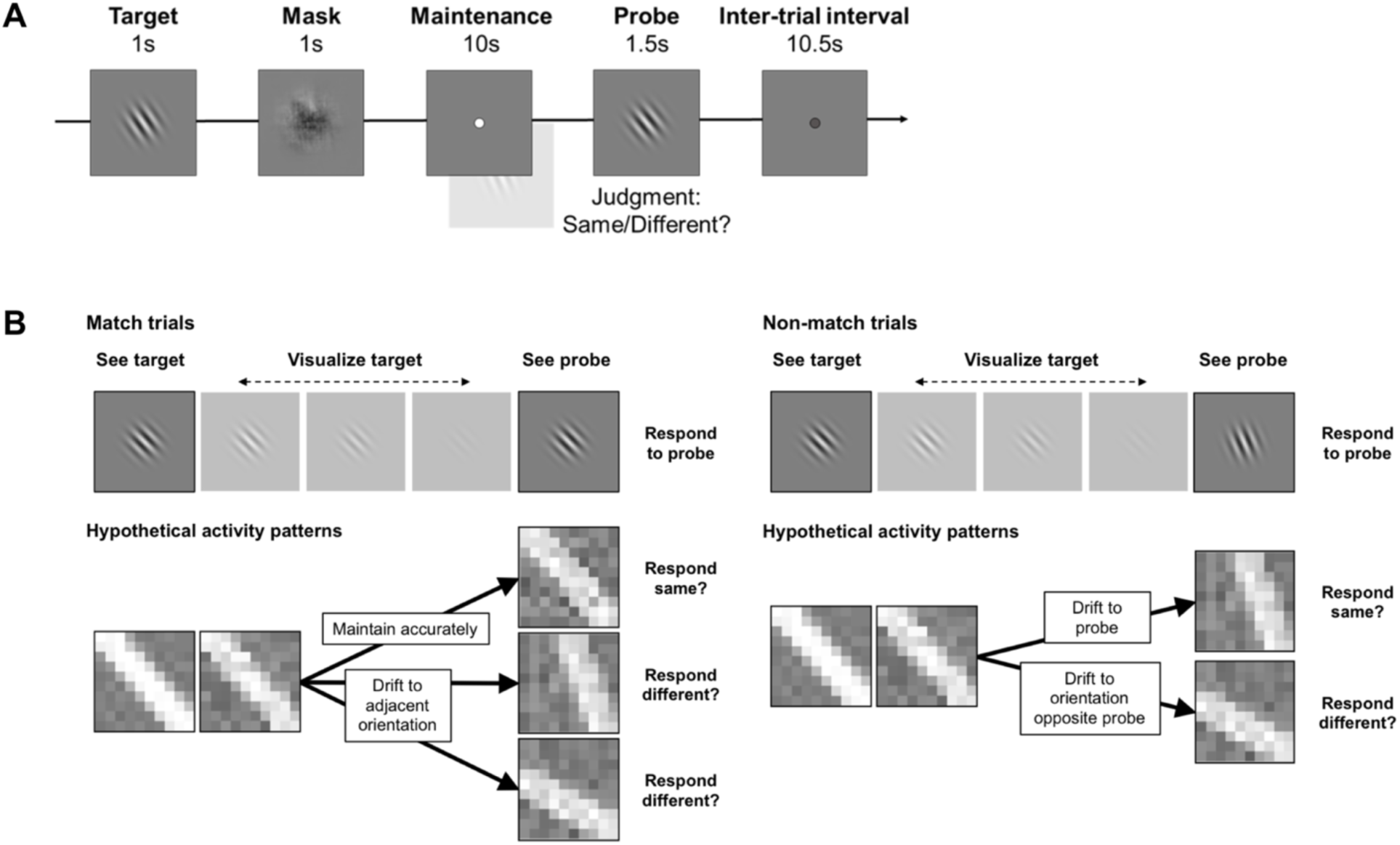
Task design and analysis approach. **(A)** Task design. Participants viewed an initial oriented Gabor patch (*target*), and held it in memory using a visualization strategy. A second patch (*probe*) then appeared, which was either the same item as the target (*match* trial), or had been rotated slightly (*non-match* trial). Participants pressed a button to indicate whether the target and probe were the *same* or *different*. Each participant only encountered a small number of discrete orientations with a fixed rotational distance between them; this distance was calibrated during a pre-scan staircasing procedure to maintain performance at approximately 75% correct. **(B)** Hypothetical brain activity patterns exemplifying *representational drift* analysis approach. For illustrative purposes, activity patterns are displayed as if they were two-dimensional images that resembled the stimuli/memoranda they represent. During target perception, patterns are presumed to be mostly veridical with little noise or directional bias. As maintenance begins, the representation persists but with decreasing signal-to-noise. The rightmost activity patterns then show hypothesized scenarios in which the patterns might drift throughout maintenance. E.g., on match trials, the pattern could remain close to the true representation (top scenario; making participants more likely to correctly report a match) or drift towards an adjacent orientation (bottom two scenarios; making participants more likely to erroneously report a non-match). Non-match trials afford another comparison that is more appropriate to that condition; activity patterns could drift towards the probe orientation (top scenario; making participants more likely to erroneously report a match) or in the opposite direction (bottom scenario; making participants more likely to correctly report a non-match).

Participants were instructed to remember the target using an imagery strategy, by visualizing it on the screen. At the end of the delay interval, a probe item (another Gabor patch) appeared for 1.5s. The probe was either identical to the target, or had been rotated by an amount specific to each participant (determined earlier; see Staircasing). Half of all trials were *match* trials, wherein the target and probe orientations were identical; the other half were *non-match* trials, wherein the probe was rotated either clockwise (50% of non-match trials) or counterclockwise from the target orientation. Participants made a *same* or *different* response by pressing one of two buttons with the index or middle fingers, respectively, of their dominant hand. Responses were only recorded if they were made while the probe was onscreen, and a short confirmation tone was played if participants responded within this timeframe, regardless of accuracy. Participants were encouraged to respond as quickly as possible while maintaining accuracy. A 10.5s fixation interval with a dark gray dot followed the probe. In the final 1s of this interval, the fixation dot changed to white to alert participants a new trial was about to begin. We chose relatively long WM delay and inter-trial intervals to allow the BOLD signal to return as much as possible to baseline levels before the probe or the next trial, respectively, thus minimizing contamination of the BOLD signals by previous events’ responses.

#### Staircasing

The first run of the DMTS task took place outside the scanner and employed a staircasing procedure. During this run, task difficulty was adjusted by varying the rotation of the probe in non-match trials according to participants’ performance, with the goal of achieving a 75% accuracy rate on the subsequent in-scanner runs. The staircasing run began with non-match probes rotated 10° relative to the target orientation. After each correct response, the non-match rotation was reduced by 1° and after each incorrect response, it was increased by 3°. After the run, a weighted average of all probe rotation values was calculated (according to an inverse exponential function over trial number, so that later trials were weighted more heavily than earlier ones), and this rotation value was used for that participant in the in-scanner runs. Probe rotation values for the in-scanner runs were capped at a maximum of 15° and a minimum of 5°, even if staircasing performance produced higher or lower values. If, during the first two runs in the scanner, a participant’s accuracy was below 60% or above 90% at the end of a run, difficulty was adjusted manually to attempt to bring performance closer to 75% on subsequent runs, and pre-adjustment runs were later removed from analysis. Due to this adjustment, one participant had two runs removed, and three participants had one run removed; all other participants completed all of their scanner runs with no difficulty adjustment needed. Task timing and procedure in the pre-scan staircasing run were largely similar to the scan runs, except that in the staircasing run, a feedback image appeared for 1s after the probe (smiley/frowny face for correct/incorrect), and the following inter-trial interval was shortened by 1s accordingly, to 9.5s. Participants received no accuracy feedback in the scanner, but did receive the confirmation tone to indicate that their response had been registered. Another difference was that during staircasing, target orientations were randomly selected from the ranges 45° ± 35° and 135° ± 35°, whereas in the scan runs, a fixed set of target orientations were used (see below).

#### Stimuli

Target and probe stimuli were large, centrally presented Gabor patches (contrast 50%, phase 0), identical for all trials and participants except for their orientations. Target orientations for each in-scanner run were drawn equally from six evenly spaced orientations, wherein the spacing was determined by the earlier staircasing run. Probe orientations were one of eight possible evenly spaced orientations; six of these were the same as the target orientations, and the last two were one additional rotation step beyond the first and last target orientations. These two extreme probe orientations occurred only once per run each, and only in non-match trials (e.g., when the most clockwise target was followed by a non-match probe rotated clockwise). In half the scan runs, targets and probes were centered around a 45° orientation; in the other half of the scan runs, targets and probes were centered around a 135° orientation (with even/odd runs alternating between the 45° and 135° base orientations; starting base orientation counterbalanced across participants). For instance, if a participant’s staircased difficulty was a step size of 10°, their six possible target orientations on a 45°-centered run would be oriented 20°, 30°, 40°, 50°, 60°, and 70°, and their eight possible probe orientations would be the same six target orientations with two additional rotations of 10° and 80°. Each 24-trial run comprised a complete and balanced set of all possible target/probe configurations for the given base orientation; four trials of each possible target position, two of which were match and two of which were non-match, and of the two non-match trials, one each in which the probe was rotated clockwise/counterclockwise. Trial orders were pseudo-randomized with the following constraints: 1) A maximum of three match or three non-match trials could occur consecutively; 2) For consecutive non-match trials, a maximum of two clockwise probe rotations, or two counterclockwise probe rotations, could occur consecutively; 3) The same target orientation was never presented in consecutive trials; 4) A previous trial’s probe orientation could not re-occur as the next trial’s target orientation (e.g., if the previous trial had used a probe orientation of 40°, the next trial’s target orientation could not have been 40°); 5) For all runs centered around the same orientation for a given participant, a sequence of the same two trials was never repeated (e.g., for all runs centered around a 45° orientation, a target orientation of 40° match trial could have been followed by a target orientation of 60° match trial only once); 6) Each trial occurred in a different position order for every run (e.g., if a given participant’s first trial in one run was a non-match trial that consisted of a target orientation at the leftmost position and probe orientation rotated clockwise, this non-match trial with the same parameters would not have been presented first in any other run).

Mask stimuli (presented immediately after encoding to reduce retinal afterimages) consisted of grayscale scene images that were randomly selected from a large set, phase-scrambled, passed through a circular Gaussian envelope matching that of the Gabor patch, rotated a random amount, and flashed at 15 Hz for the duration of the 1s mask interval.

### Retinotopic mapping task

Following the DMTS task runs, participants completed four ∼2.5-minute runs of a standard retinotopic mapping task. Participants viewed a wedge-shaped checkerboard pattern that rotated about a central fixation dot at 2.5 cycles per minute and flickered at a rate of 10 reversals per second. The wedge rotated either clockwise or counterclockwise for the entire run, alternating between runs. Participants were instructed to maintain fixation while watching for a brief color change in the fixation dot, which occurred approximately every 10 seconds on average. Whenever they detected this change, they pressed a button with the index finger on their dominant hand.

### fMRI data acquisition

Scanning was performed on a Siemens 3T Skyra system with a 32-channel head coil. Functional scans used a multiband echoplanar imaging sequence (Feinberg et al. 2010; Moeller et al. 2010) with TR=1000ms, TE=30ms, 100−100 in-plane resolution, 60 axial slices with thickness 2.2mm and 0mm skip, field of view = 220mm (overall voxel size: 2.2 × 2.2 × 2.2mm), flip angle = 60°, interleaved acquisition with multiband factor of 4. Scans were prescribed with slices parallel to the anterior commissure/posterior commissure line and positioned for whole-brain coverage. Participants performed six runs of the main DMTS task with 580 volumes (9 minutes, 40 seconds) per run and four runs of the retinotopic mapping task with 160 volumes (2 minutes, 40 seconds) per run. In each run, to allow the fMRI signal to reach steady-state before onset of the first trial, the scanner ran for 4 seconds/volumes without collecting data, and an additional 4 seconds/volumes of collected data were discarded from the beginning of each run. T1-weighted MPRAGE anatomical images were also collected for each participant at the beginning of each scan session (TR=2200ms, TE=3.37ms, 256 × 256 × 192 1mm-isotropic voxels, sagittal slice prescription).

One participant completed only five DMTS runs due to technical difficulties; another completed only half of her final DMTS run before it was aborted due to physical discomfort in the scanner. (In addition, as noted above, one additional participant had two runs removed and three participants had one run removed due to manual difficulty adjustments on early scan runs.) All participants completed all four runs of the retinotopy task.

### fMRI data preprocessing

Initial processing of fMRI data was performed using SPM8 (Statistical Parametric Mapping; Wellcome Trust Centre for Neuroimaging, University College London, UK). Data were motion-corrected, and all of a participant’s functional runs were coregistered to a mean image of that participant’s first functional run after motion correction. Each participant’s T1 anatomical image was then coregistered to the Montreal Neurological Institute (MNI) average structural template image. Participants’ motion-corrected functional images were then coregistered to this anatomical image, and all functional images were resampled. Thus, all participant data were approximately aligned to MNI space but only affine transformations were applied, keeping data in individual-subject space with no nonlinear warping and only a single resampling step at the end. For the DMTS task, fMRI signal values at each timepoint were then z-scored across the entire volume to control for signal fluctuations over time. These z-scored versions of the functional volumes were used as the basis of all pattern analyses.

### Omnibus visual cortex ROI definition

Based on the knowledge that multiple retinotopic visual areas represent information about items held in visual WM (Harrison and Tong 2009), our main analyses were based on a functionally defined ROI comprising the most responsive retinotopically mapped voxels in the brain, irrespective of their anatomical location. Data from the retinotopic mapping task were used to identify the 1000 voxels that responded most robustly to the rotating checkerboard. Each voxel’s timecourse during the retinotopy task was Fourier transformed, and the amplitude at the frequency corresponding to the rotation of the checkerboard wedge was extracted. This amplitude was converted to a Pearson correlation r-value, Fisher z′-transformed, and averaged across the four retinotopic mapping runs. The voxels with the highest mean z′-transformed correlation values across all four mapping runs were selected. Then, in the main DMTS runs, this set of 1000 voxels was extracted from each z-scored volume of functional data and used for all subsequent pattern analyses.

### Visual areas V1, V2, and V3 ROI definition

To supplement the primary ROI analyses and determine whether there were meaningful differences in results between retinotopic visual regions, we also repeated our main analyses for areas V1–3 individually. For each participant, probabilistic maps of V1–3 were obtained using an anatomical template of the cortical surface with population-defined maps of those visual areas overlaid (Benson et al. 2012). This method has been shown to have accuracy comparable to manually defining these ROIs for each participant (Benson et al. 2012, 2014) and eliminates the possibility of experimenter bias in ROI definition. For each participant, we used FreeSurfer 5.3 (Reuter et al. 2010) to reconstruct cortical surface maps from the T1 anatomical image, calculate the transformation from the surface template to the participant’s cortical surface, apply the V1–3 template maps, and then re-transform those subject-specific V1–3 surface maps back into 3-D volume space aligned with their pre-processed functional images. Then, we masked the results of the retinotopy analysis above with our V1–3 ROI definitions and selected the 500 most responsive retinotopically mapped voxels from each area. For each ROI, voxel patterns during the DMTS task were extracted as for the main visual cortex ROI above.

### fMRI pattern similarity and representational drift

We calculated pattern similarity values and *representational drift* indices in our four ROIs described above to determine how ongoing changes in brain activity patterns corresponded with performance. Separate analyses were conducted for match trials (where target and probe orientations were the same) and non-match trials (where target and probe orientations were different). Statistical comparisons focused on the differences in pattern similarity or drift index between accurate and inaccurate trials during timepoints representing the encoding, maintenance, and probe intervals of the DMTS task (see Results).

#### Prototypical activity patterns

For each participant and each ROI, we first obtained prototypical activity patterns for each unique target orientation seen during the DMTS task. These patterns represent the “canonical” version of the expected activity patterns corresponding to visual processing of each task-relevant orientation; pattern similarity during WM to these prototypical patterns should thus reflect successful re-instantiation of those orientations. The prototypical voxel patterns for each orientation were calculated by averaging voxel patterns from all trials in which that orientation was the target, using patterns from fMRI volume 5 within each trial (with TR=1000ms, this volume represented the peak activity occurring in response to the target presentation at t=0s of the trial, after accounting for BOLD response delay). We thus obtained prototypical activity patterns for twelve unique orientations in total (e.g., the hypothetical participant with a staircased difficulty step size of 10° would have prototypical activity patterns for the twelve orientations 20°–70° and 110°–160°, inclusive, in steps of 10°). Because target and probe stimuli used the same orientations (with the exception of the two most extreme probe orientations), prototypical activity patterns could therefore be obtained for almost all task-relevant orientations in a scan session. Trials in which the participant did not respond (only 3.5% of all trials) were still used to generate prototypical patterns but not used in any subsequent analyses based on accuracy.

#### Pattern similarity and representational drift calculations

Several task-relevant orientations and their corresponding voxel patterns formed the basis of our calculations. In addition to the *target orientation* and the *probe orientation* (note that the probe orientation was the same as the target on match trials but a different, adjacent orientation on non-match trials), we also defined two *target-adjacent* control orientations for match trials and a *probe-opposite* control orientation for non-match trials. The target-adjacent orientations were the two possible stimulus orientations adjacent to the target on a given trial (one rotated one step clockwise, the other one step counterclockwise). The probe-opposite orientation was the orientation adjacent to the target on a given trial that was not the probe (i.e., rotated one step away from the target in the opposite direction from the probe).

Raw pattern similarity values were calculated by taking the Euclidean distance between two vectors, one representing the voxel pattern in a given ROI at a specific timepoint and the other a prototypical activity pattern in that ROI for one of the task-relevant orientations. For each trial, raw pattern similarity values were calculated for every fMRI volume (1–24). To account for any differences in initial representation, values at each timepoint were subtracted from the value at fMRI volume 1. Thus, all pattern similarity timelines began at 0 and the sign was inverted so that positive values indicate higher similarity (lower distance). As the TR was 1000ms, volume 1 was collected between t=0s and t=1s, volume 2 between t=1s and t=2s, and so on; in all figures, fMRI volumes are represented by the average time of their collection (0.5s, 1.5s, etc.). In all analyses, pattern similarity indices were calculated separately for accurate and inaccurate trials, and all individual-trial timelines were averaged within participants before entering them into statistical analyses.

For match trials, we calculated raw pattern similarity timelines comparing ongoing activity to each of the following prototypical activity patterns: the *target pattern* (**Figure 2A**; i.e., that trial’s target orientation) and the two *target-adjacent* control orientations defined above. We then averaged those target-adjacent pattern similarities to obtain a single combined measure for the control orientations (**Figure 2B**). Two participants were removed from the match trial analysis because they had very few inaccurate match trials.

**Figure 2.**
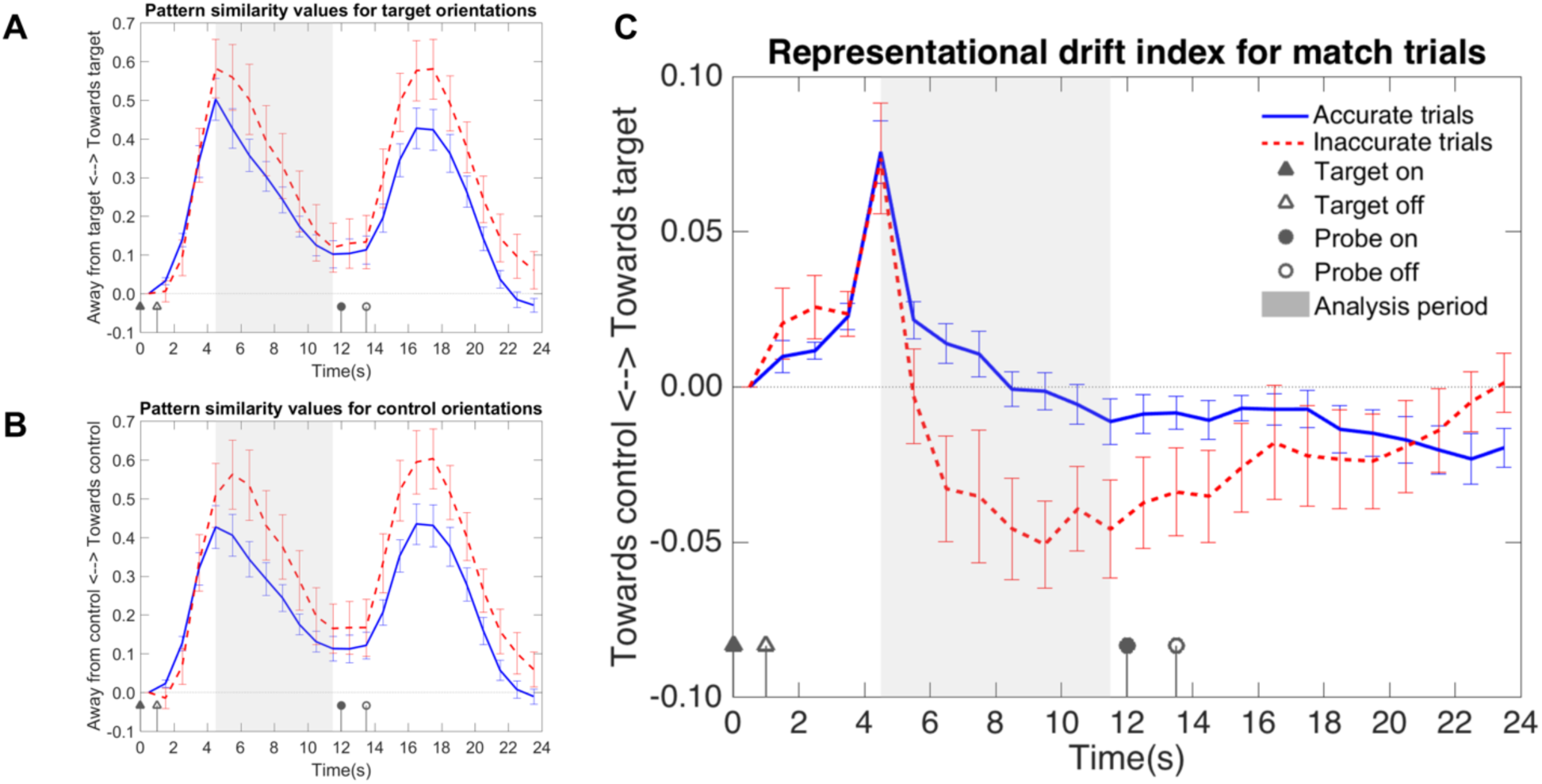
Match trials: Pattern similarity and representational drift in visual cortex ROI. Timelines for pattern similarity and representational drift for match trials in visual cortex ROI. Accurate and inaccurate trials are plotted separately. All plots depict the timecourse of an entire 24-second trial (fMRI volumes 1–24). Data are represented as mean ± SEM. To examine effects occurring during the maintenance period, we analyzed an eight-second time window (shaded in gray), starting from the BOLD peak of target encoding (fMRI volume 5) and ending just before the probe appeared onscreen (volume 12). Note that the legend shown in (C) applies to all three plots. **(A) Match trials: Target pattern similarity.** Pattern similarity between each timepoint of the trial and the prototypical activity pattern for the target orientation. Positive values indicate more similar activity patterns to the prototypical target pattern for that trial, while negative values indicate less pattern similarity to the prototypical target pattern for that trial. There was no difference between accurate and inaccurate trials during the maintenance period, but pattern similarity was higher for inaccurate trials at the timepoint representing the BOLD peak of probe presentation (fMRI volume 17, t=16.5s). **(B) Match trials: Control orientation pattern similarity.** Pattern similarity between each timepoint of the trial and the prototypical activity patterns for target-adjacent control orientations, which were the two orientations adjacent to the target. (Pattern similarity values from each control orientation were averaged to produce a single timeline.) Positive values indicate more similar activity patterns to prototypical target-adjacent orientations for that trial, while negative values indicate less pattern similarity to prototypical target-adjacent orientations for that trial. Mirroring the target pattern similarity analysis in (A), there was no difference between accurate and inaccurate trials during the maintenance period, but pattern similarity was higher for inaccurate trials at the timepoint representing the BOLD peak of probe presentation. **(C) Match trials: Representational drift.** Representational drift index for each timepoint of the trial. We subtracted the target-adjacent pattern similarities from the target pattern similarities to create a single index of *representational drift*, which conveys whether any changes in brain activity patterns represented a net drift in representational similarity towards either the target orientation or an adjacent (competing) orientation. Positive values indicate representational drift towards the target orientation for that trial, while negative values indicate representational drift towards target-adjacent orientations for that trial. We found a significant main effect of Accuracy during maintenance as well as a significant quadratic effect of Time by Accuracy; net representational drift away from the target on inaccurate trials was greater in the middle portion of the maintenance period than at the beginning or the end.

For non-match trials, we calculated raw pattern similarity timelines comparing ongoing activity to each of the following prototypical activity patterns: the *target pattern* (**Figure 3A**; same as for match trials), the *probe pattern* (**Figure 3B**; i.e., that trial’s non-matching probe orientation), and the *probe-opposite* control orientation defined above (**Figure 3C**).

**Figure 3.**
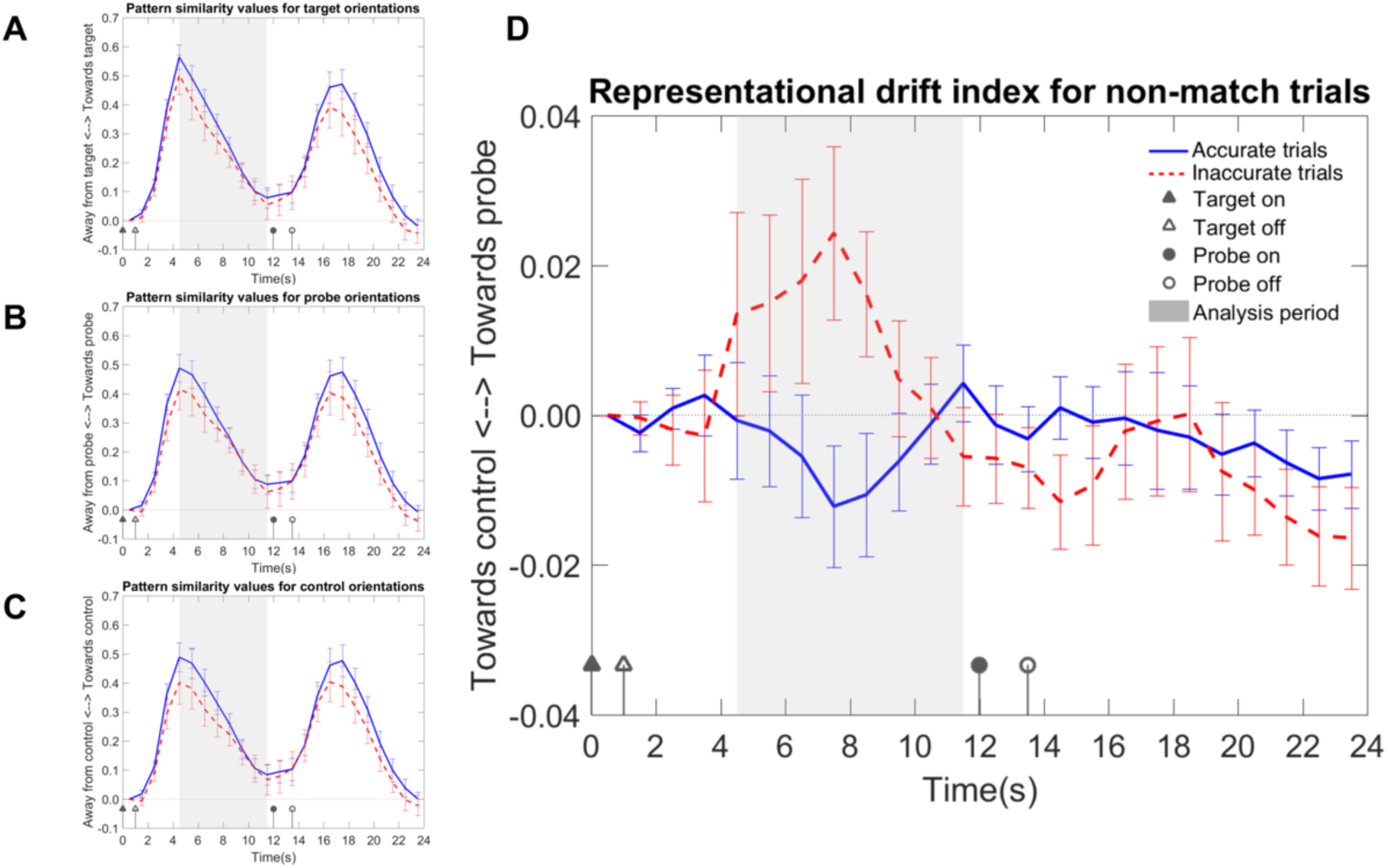
Non-match trials: Pattern similarity and representational drift in visual cortex ROI. Timelines for pattern similarity and representational drift for non-match trials in visual cortex ROI. Accurate and inaccurate trials are plotted separately. All plots depict the timecourse of an entire 24-second trial (fMRI volumes 1–24). Data are represented as mean ± SEM. As in match trials, to examine effects occurring during the maintenance period, we analyzed an eight-second time window (shaded in gray), starting from the BOLD peak of target encoding (fMRI volume 5) and ending just before the probe appeared onscreen (volume 12). Note that the legend shown in (D) applies to all four plots. **(A) Non-match trials: Target pattern similarity.** Pattern similarity between each timepoint of the trial and the prototypical activity pattern for the target orientation. Positive values indicate more similar activity patterns to the prototypical target pattern for that trial, while negative values indicate less pattern similarity to the prototypical target pattern for that trial. Pattern similarity was initially numerically higher for accurate trials than inaccurate, but the difference disappeared by the end of the maintenance period. **(B) Non-match trials: Probe pattern similarity.** Pattern similarity between each timepoint of the trial and the prototypical activity pattern for the probe orientation. Positive values indicate more similar activity patterns to the prototypical probe pattern for that trial, while negative values indicate less pattern similarity to the prototypical probe pattern for that trial. Similar to target pattern similarity in (A), probe pattern similarity was initially numerically higher for accurate trials than inaccurate, but the difference disappeared by the end of the maintenance period. **(C) Non-match trials: Control pattern similarity.** Pattern similarity between each timepoint of the trial and the prototypical activity pattern for the probe-opposite control orientation, which was the orientation that, like the probe, was adjacent to the target orientation, but was rotated in the opposite direction. Positive values indicate more similar activity patterns to the prototypical control orientation pattern for that trial, while negative values indicate less pattern similarity to the prototypical control orientation pattern for that trial. Similar to target and probe pattern similarity (panels A and B, respectively), control orientation pattern similarity was initially numerically higher for accurate trials than inaccurate, but the difference disappeared by the end of the maintenance period. **(D) Non-match trials: Representational drift.** Representational drift index for each timepoint of the trial. We subtracted the control pattern similarities from the probe pattern similarities to create a single index of *representational drift*, which conveys whether any changes in brain activity patterns represented a net drift in representational similarity towards either the probe orientation or the probe-opposite control orientation. Positive values indicate representational drift towards the probe orientation for that trial, while negative values indicate representational drift towards the probe-opposite orientation for that trial. We found a significant quadratic effect of Time by Accuracy; inaccurate trials showed net representational drift towards the probe in the middle portion of the maintenance period, but not at the beginning or end.

However, raw pattern similarity values on their own are an insufficient means of indicating the fidelity of participants’ WM representations of a *specific* item, as it is possible for brain activity patterns to change in similarity towards or away from multiple representations at once. For example, if a participant became distracted and stopped paying attention mid-trial, his/her brain activity patterns would likely become less similar to all task-relevant orientations at the same time. Thus, we also calculated *representational drift* indices that, unlike raw pattern similarity values, are capable of conveying whether activity patterns are drifting more towards one particular representation than another.

Representational drift indices were calculated by subtracting one raw pattern similarity timeline from another, allowing a direct comparison between the two. As all raw pattern similarity timelines were baselined to begin at 0, an advantage of this approach is that the representational drift index is guaranteed to correspond to a net change in pattern similarity towards a specific representation since the beginning of the trial, with positive values indicating net drift towards one orientation and negative values indicating net drift towards the other.

For match trials, we calculated representational drift timelines comparing the target orientation to the target-adjacent control orientations (**Figure 2C**, which represents a subtraction of the raw pattern similarity values in **Figure 2B** from those in **Figure 2A**; i.e., PS_target_ – PS_control_). Thus, positive values indicate changes in pattern similarity towards the target orientation, while negative values indicate changes in pattern similarity towards the control orientations. Because any general effects such as distraction should affect both raw pattern similarity timelines equally, subtracting those timelines should cancel out such non-specific influences on brain activity patterns, and any effects seen in the representational drift index should be due to differences in particular WM representations.

For non-match trials, we calculated representational drift timelines comparing the probe orientation to the probe-opposite control orientation (**Figure 3D**, which represents a subtraction of the values in **Figure 3C** from those in **Figure 3B**; i.e., P_Sprobe_ – P_Scontrol_). Thus, positive values indicate changes in pattern similarity towards the orientation that will ultimately be probed, while negative values indicate changes in pattern similarity towards the control orientation.

As no prototypical activity patterns existed for the most extreme probe orientations (because those orientations never were seen as targets), raw pattern similarities and drift indices based on those orientations could not be calculated. Thus, we did not analyze either match or non-match trials where the target orientation was one of the endpoints of the range of possible target orientations for that run (e.g., in a target set using orientations of 20°, 30°, 40°, 50°, 60°, and 70°, the analysis would exclude trials where the target was 20° or 70°), as those calculations would have required non-existent prototypical activity patterns for either the probe or control orientations.

## RESULTS

We calculated *representational drift* in fMRI activity patterns during the delay period of a delayed match-to-sample WM task to determine how ongoing changes in the quality of activity pattern representations correspond with performance. Participants viewed an initial oriented Gabor patch (the *target*), held it in memory for an 11-second maintenance period, and then saw a second Gabor patch (the *probe*) that was either the same orientation as the target (*match* trial) or rotated slightly (*non-match* trial). Participants responded with a button press to indicate whether they thought the target/probe orientations were the same or different (see **Figure 1A**). Each participant only encountered a small number of discrete orientations with a fixed rotational distance between them; this distance was calibrated during a pre-scan staircasing procedure (for details, see Methods) to maintain performance at approximately 75% correct. Indices of representational drift were calculated based on changes in multi-voxel pattern similarity during maintenance; these indicated whether brain activity patterns drifted toward (become more similar to) or away from (became less similar to) the *prototypical* (average) activity pattern associated with visual perception of a given orientation (e.g., the target or the probe; see **Figure 1B** for an illustrative example). The timecourses of these representational drift indices were then compared between accurate and inaccurate trials to determine how representational drift in brain activity related to task performance.

### Behavioral performance

Each run comprised 24 trials, 12 *match* trials and 12 *non-match* trials. Each trial lasted 24s (see **Figure 1A**) and thus each run was approximately 10 minutes long. All participants completed 4– 6 task runs. Mean accuracy across all runs was 72.0% (SD = 7.4%) and mean response time was 921 ms (SD = 94 ms). 3.5% of trials received no response; the average number of trials completed by each participant and included in our analyses was 132.

### fMRI analysis: Omnibus visual cortex region of interest (ROI)

For each participant, we identified an omnibus visual cortex ROI comprising the top 1000 visually responsive voxels in the brain, based on which voxels activated most in a standard retinotopic mapping task that followed the main task. For this ROI, we first report basic “raw” pattern similarity values at each point in the trial between that timepoint’s activity pattern and the prototypical activity patterns (the average activity patterns corresponding to visual processing of that orientation; see Methods for details) for the critical orientations used in that trial. Temporal resolution was 1s. To examine effects at encoding (t=0s in the trial) and probe presentation (t=12s), we allowed 4–5s for blood-oxygenation-level-dependent (BOLD) signal delay and ran paired t-tests comparing pattern similarity between accurate and inaccurate trials at fMRI volumes 5 and 17 of the trial, respectively. To examine effects occurring during the maintenance period, we ran a linear repeated-measures analysis of variance including factors for the main effect of Accuracy and the linear and quadratic components of the Time x Accuracy interaction. This analysis was run on an eight-second time window, starting from the BOLD peak of target encoding (fMRI volume 5) and ending just before the probe appeared onscreen (volume 12). All plots (**Figures 2–4**) depict the timecourse of an entire 24-second trial (fMRI volumes 1–24).

#### Match trials: Target pattern similarity

**Figure 2A** shows, for *match* trials, pattern similarity between each timepoint of the trial and the prototypical activity pattern for the *target* orientation. A paired t-test between accurate and inaccurate trials was not significant at encoding (p = .237), but was at probe (t(17) = 2.14, p = .047). Specifically, at probe, pattern similarity was greater on inaccurate than accurate trials. This suggests that, counterintuitively, when there was more pattern similarity between that trial’s activity pattern at probe and the prototypical target activity pattern, participants were more likely to (incorrectly) report a non-match. Our analysis of the maintenance period showed no significant effect of Accuracy (p = .241), and no linear (p = .194) or quadratic (p = .286) trends for the Time x Accuracy interaction. This suggests that the raw pattern similarity between the prototypical target representation and participants’ activity patterns during maintenance did not have a measurable effect on task accuracy for *match* trials.

#### Match trials: Control orientation pattern similarity

We then calculated pattern similarity between each timepoint of the trial and the prototypical patterns for two *control orientations*, which were the two orientations adjacent to the target. This allowed us to determine to what extent any pattern similarity effects seen in **Figure 2A** were unique to the target orientation, or conversely, whether they were due to less specific phenomena (e.g., general inattention on some trials, leading to inaccurate responses). **Figure 2B** shows, for *match* trials, pattern similarity between each timepoint of the trial and the prototypical activity patterns for the control orientations. Similar to the analysis of target pattern similarity, a paired t-test between accurate and inaccurate trials was not significant at encoding (p = .237), but was at probe (t(17) = 2.27, p = .037). Pattern similarity at probe was again greater on inaccurate than accurate trials, suggesting that greater pattern similarity to any task-relevant orientation (target or target-adjacent) at retrieval is associated with (incorrect) reporting of a non-match. Our analysis of the maintenance period, as for target orientations, showed no significant effect of Accuracy (p = .093), and no linear (p = .340) or quadratic (p = .096) trends for the Time x Accuracy interaction. Thus, the raw pattern similarity during maintenance did not appear to have a measurable effect on *match* trial accuracy for either the target orientation or our target-adjacent control orientations.

#### Match trials: Representational drift

The pattern similarity analyses above did not strongly support any effects of raw pattern similarity during maintenance on participants’ accuracy, for either the target orientation or the target-adjacent control orientations. Furthermore, the similarity between the timelines shown in **Figures 2A–B** suggests that raw pattern similarity alone may not be a good indicator of the quality of participants’ WM representations for the target orientation, specifically. However, we hypothesized that the *difference* in raw pattern similarities between the target and control orientations (PS_target_ – PS_control_) might better reflect the quality of the target’s WM representation. Because any general effects such as distraction should affect both the target and control pattern similarity timelines equally, subtracting those timelines should cancel out such non-specific influences on brain activity patterns. Thus, we subtracted the target-adjacent pattern similarities from the target pattern similarities to create a single index of *representational drift*. This index conveys, independent of any more general factors (e.g., waxing and waning attention), a true shift in pattern space towards either the target orientation or the control orientations; i.e., whether any changes in brain activity patterns represented a net drift in representational similarity towards either the target orientation or an adjacent (competing) orientation.

**Figure 2C** shows, for *match* trials, the representational drift index for each timepoint of the trial. A paired t-test between accurate and inaccurate trials was not significant at either encoding (p = .866) or probe (p = .622). However, our analysis of the maintenance period showed a significant main effect of Accuracy (F(1,17) = 5.84, p = .027), with more net representational drift away from the target orientation (towards target-adjacent orientations) on inaccurate trials. The Time x Accuracy interaction showed no significant linear trend (p = .232) but did show a significant quadratic trend (F(1,17) = 5.52, p =.031), where net representational drift away from the target on inaccurate trials was greater in the middle portion of the maintenance period than at the beginning or at the end. This suggests that participants were more likely to incorrectly report a non-match when their activity patterns drifted away from the target orientation and towards target-adjacent orientations; furthermore, this effect was largest in the middle portion of the maintenance period.

#### Non-match trials: Target pattern similarity

**Figure 3A** shows, for *non-match* trials, pattern similarity between each timepoint of the trial and the prototypical activity pattern for the target orientation. A paired t-test between accurate and inaccurate trials was not significant at either encoding (p = .173) or probe (p = .132). Our analysis of the maintenance period showed no significant effect of Accuracy (p = .281) and no quadratic trend (p = .863) for the Time x Accuracy interaction. However, there was a near-significant linear trend for the interaction (F(1,19) = 4.17, p = .055) where pattern similarity was initially numerically higher for accurate trials than inaccurate, but the difference between accurate and inaccurate trials disappeared by the end of the maintenance period. This suggests that when there was more initial pattern similarity between that trial’s activity pattern and the prototypical target activity pattern, participants may have been more likely to (correctly) report a non-match, even though such starting differences were negated later in the maintenance period.

#### Non-match trials: Probe pattern similarity

**Figure 3B** shows, for *non-match* trials, pattern similarity between each timepoint of the trial and the prototypical activity pattern for the probe orientation. Similar to the target pattern similarity analysis above, a paired t-test between accurate and inaccurate trials was not significant at either encoding (p = .103) or probe (p = .199). Also closely mirroring the analysis of target pattern similarity, our analysis of the maintenance period showed no significant effect of Accuracy (p = .362), and no quadratic trend (p = .329) for the Time x Accuracy interaction, but there was a significant linear trend for the Time x Accuracy interaction (F(1,19) = 4.82, p = .041). As above, pattern similarity during maintenance was initially numerically higher for accurate trials than inaccurate, but that difference disappeared by the end of the maintenance period.

#### Non-match trials: Control orientation pattern similarity

We also calculated pattern similarity on *non-match* trials for a control orientation that, like the probe, was adjacent to the target orientation, but was rotated in the opposite direction (the *probe-opposite* orientation). As with the control orientation analysis for match trials, this allowed us to determine to what extent any pattern similarity effects seen in **Figures 3A–B** were unique to the target/probe orientations, or conversely, whether they were due to less specific phenomena. **Figure 3C** shows pattern similarity between each timepoint of the trial and the prototypical activity pattern for the probe-opposite control orientation. A paired t-test between accurate and inaccurate trials trended towards significance at encoding (t(19) = 2.01, p = .059) but not at probe (p = .172). Specifically, at encoding, pattern similarity was greater on accurate than inaccurate trials. Similar to the analyses of target and probe pattern similarity, our analysis of the maintenance period showed no significant main effect of Accuracy (p = .159), and no quadratic trend (p = .922) for the Time x Accuracy interaction, but a significant linear trend for the interaction (F(1,19) = 7.70, p = .012) where pattern similarity was initially numerically higher for accurate trials than inaccurate, but the difference disappeared by the end of the maintenance period. This suggests that greater pattern similarity to any task-relevant orientation (target, probe, or control) at encoding is associated with (correct) reporting of a non-match. However, none of these analyses of raw pattern similarity appeared to suggest that the fidelity of participants’ WM representations for specific orientations during the maintenance period had an effect on accuracy.

#### Non-match trials: Representational drift

Given that the raw pattern similarity analyses above did not strongly support any orientation-specific effects on participants’ accuracy, as well as the general similarity between the timelines shown in **Figures 3A–C**, it appeared that raw pattern similarity alone (as for *match* trials) may not be a good indicator of the quality of participants’ WM representations for specific orientations. However, similar to our representational drift analysis for *match* trials, we hypothesized that the *difference* in raw pattern similarities between the probe and control orientations (PS_probe_ – PS_control_) might better convey consequential changes in participants’ WM representations during maintenance. Thus, we subtracted the control pattern similarities from the probe pattern similarities to create a single index of *representational drift*. Critically, this index of representational drift during non-match trials, unlike the representational drift index for match trials above, is capable of reflecting a directional effect of representational drift on accuracy; in other words, this index can indicate whether participants are more likely to (incorrectly) report a match when their WM representations drift towards the non-matching probe orientation than when their WM representations drift away from the probe orientation and towards the probe-opposite control orientation.

**Figure 3D** shows, for *non-match* trials, the representational drift index for each timepoint of the trial. A paired t-test between accurate and inaccurate trials was not significant at either encoding (p = .307) or probe (p = .896). Our analysis of the maintenance period showed no significant main effect of Accuracy (p = .206) and no linear trend (p = .191) for the Time x Accuracy interaction. However, there was a significant quadratic trend for the interaction (F(1,19) = 6.66, p = .018), with inaccurate trials having net representational drift towards the probe in the middle portion of the maintenance period, but not at the beginning or end. This suggests that participants were more likely to incorrectly report a match when their activity patterns drifted towards the probe orientation (or, conversely, more likely to correctly report a non-match when their activity patterns drifted towards the probe-opposite orientation), with the representational drift effect being maximal in the middle portion of the maintenance period.

### fMRI analysis: Visual areas V1, V2, and V3

Because past research has found some differences between individual visual areas during WM maintenance (Harrison and Tong 2009), we also examined visual areas V1, V2, and V3 separately. For each participant, we obtained probabilistic maps of areas V1–3 using an anatomical template of the cortical surface (Benson et al. 2012). Within each of those areas, we then identified an ROI comprising the top 500 visually responsive voxels (based on our standard retinotopic mapping task) in that area. For each of these ROIs, we report the same *representational drift* indices described in the omnibus visual cortex ROI analyses (**Figures 2C** and **3D**). For *match* trials, this index reflects a net drift in representational similarity towards either the target orientation or an adjacent (competing) orientation; for *non-match* trials, this index reflects a net drift in representational similarity towards either the probe orientation or the probe-opposite control orientation.

As above, we ran paired t-tests between accurate and inaccurate trials at encoding (fMRI volume 5 in the trial) and probe (fMRI volume 17), as well as linear repeated-measures analyses of variance for the maintenance interval (fMRI volumes 5–12), including factors for the main effect of Accuracy and the linear and quadratic components of the Time x Accuracy interaction.

#### Match trials: Representational drift

**Figure 4A** shows, for *match* trials, the representational drift index for each timepoint of the trial. A paired t-test between accurate and inaccurate trials was not significant for any visual area at encoding (V1: p = .423; V2: p = .860; V3: p = .877) or at probe (V1: p = .719; V2: p = .633; V3: p = .895). Our analysis of the maintenance period showed no main effect of Accuracy for any visual area (V1: p = .393; V2: p = .558; V3: p = .198), and no linear trend for the Time x Accuracy interaction (V1: p = .457; V2: p = .891; V3: p = .376). Areas V2 and V3 also showed no quadratic trend for the interaction (V2: p = .143; V3: p = .097). However, area V1 showed a significant quadratic trend (F(1,17) = 6.22, p = .023), with inaccurate trials having net representational drift away from the target in the middle portion of the maintenance period, but not at the beginning or end. As in the omnibus visual cortex ROI, this suggests that participants were more likely to (incorrectly) report a non-match when their activity patterns in V1 drifted away from the target orientation and towards target-adjacent orientations; furthermore, this effect primarily occurred in the middle portion of the maintenance period.

**Figure 4.**
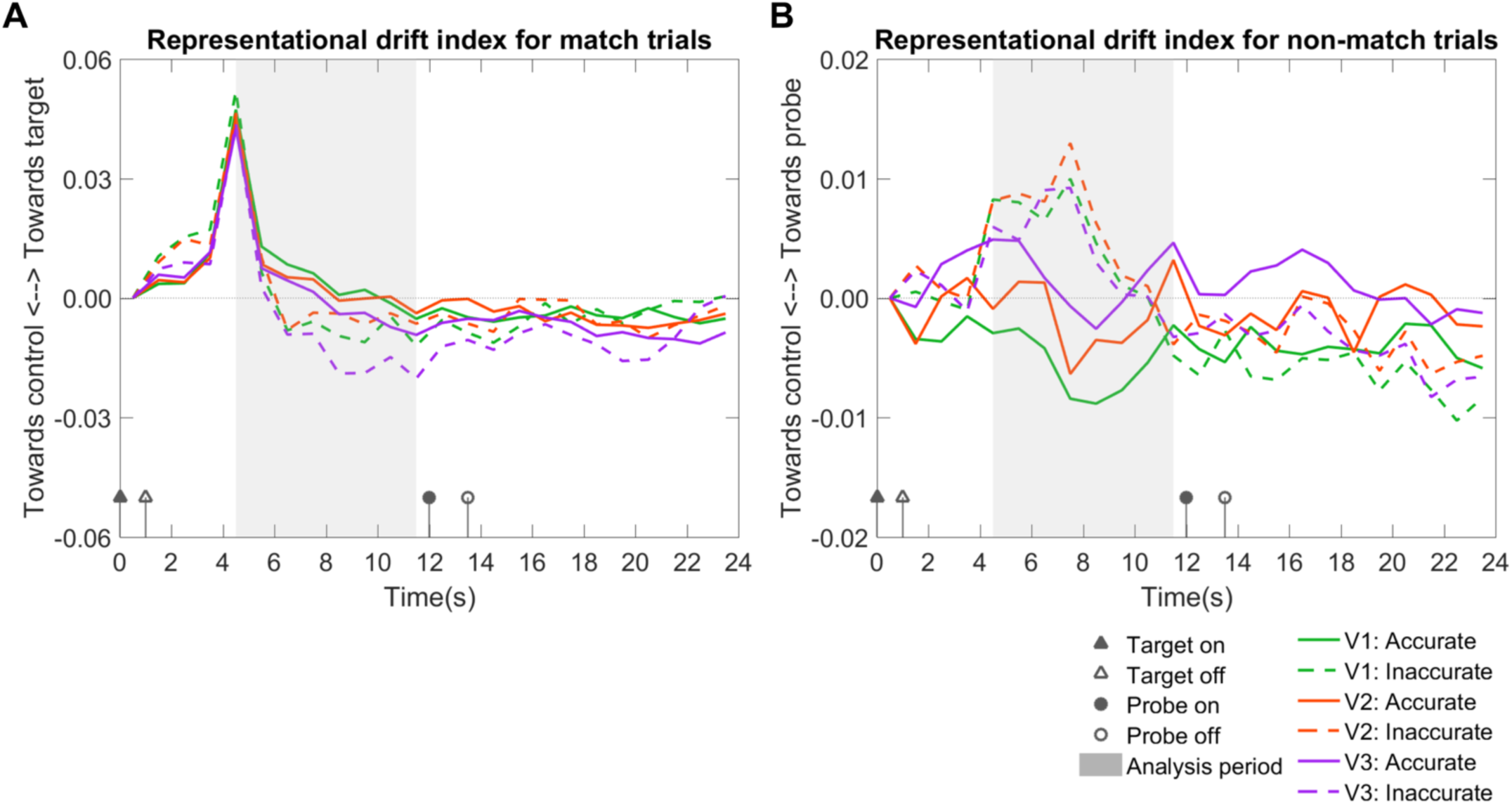
Representational drift for match and non-match trials in visual areas V1, V2, and V3. For each of these ROIs, we report the same representational drift indices described in the omnibus visual cortex ROI analyses (Figures 2C and 3D). Each ROI is plotted on a separate line. Accurate and inaccurate trials are plotted separately. All plots depict the timecourse of an entire 24-second trial (fMRI volumes 1–24). As for the previous analyses, we analyzed an eight-second time window (shaded in gray), starting from the BOLD peak of target encoding (fMRI volume 5) and ending just before the probe appeared onscreen (volume 12). Note that the legends apply to both panels. **(A) Match trials: Representational drift.** Representational drift index for each timepoint of the trial. This was calculated using the same procedure as for the omnibus visual cortex ROI (Figure 2C). Positive values indicate representational drift towards the target orientation for that trial, while negative values indicate representational drift towards target-adjacent control orientations for that trial. For areas V2 and V3, there was no significant difference between accurate and inaccurate trials during maintenance. In area V1, we found a significant quadratic effect of Time by Accuracy; net representational drift away from the target on inaccurate trials was greater in the middle portion of the maintenance period than at the beginning or the end, generally mirroring the omnibus visual cortex ROI analysis. **(B) Non-match trials: Representational drift.** Representational drift index for each timepoint of the trial. This was calculated using the same procedure as for the omnibus visual cortex ROI (Figure 3D). Positive values indicate representational drift towards the probe orientation for that trial, while negative values indicate representational drift towards the probe-opposite control orientation for that trial. For areas V1 and V2, we found significant (V1) or near-significant (V2) main effects of Accuracy during maintenance, with net representational drift towards the probe on inaccurate trials but towards the control orientation on accurate trials. In areas V1 and V3, we found significant (V1) or near-significant (V3) quadratic effects of Time by Accuracy; representational drift towards the probe was greater in the middle portion of the maintenance period than at the beginning or end for inaccurate trials, and vice versa for accurate trials, which generally mirrored the omnibus visual cortex ROI analysis.

#### Non-match trials: Representational drift

**Figure 4B** shows, for *non-match* trials, the representational drift index for each timepoint of the trial. A paired t-test between accurate and inaccurate trials was not significant for any visual area at encoding (V1: p = .117; V2: p = .102; V3: p = .507) or at probe (V1: p = .954; V2: p = .948; V3: p = .648). Our analysis of the maintenance period showed no significant main effect of Accuracy for area V3 (p = .655). However, there was a significant main effect of Accuracy for area V1 (F(1,19) = 5.35, p = .032) and a near-significant effect of Accuracy for area V2 (F(1,19) = 3.75, p = .068), with net representational drift towards the probe on inaccurate trials but towards the control orientation on accurate trials. This suggests participants were more likely to (incorrectly) report a match when their activity patterns drifted towards the probe orientation (and more likely to correctly report a non-match when their activity patterns drifted towards the probe-opposite control orientation). There was no significant linear trend for the Time x Accuracy interaction in any visual area (V1: p = .184; V2: p = .162; V3: p = .277), and no quadratic trend for the interaction in area V2 (p = .094). However, there was a significant quadratic trend for the interaction in area V1 (F(1,19) = 5.04, p = .037) and a near-significant effect in area V3 (F(1,19) = 4.11, p = .057), where representational drift towards the probe was greater in the middle portion of the maintenance period than at the beginning or at the end for inaccurate trials, and vice versa for accurate trials.

Overall, for both match and non-match trials, the results of these analyses performed on areas V1–3 qualitatively mirror those found in the omnibus visual cortex ROI, although the effects were only statistically significant in V1 and not in V2 or V3.

## DISCUSSION

We calculated *representational drift* in fMRI activity patterns during the delay period of a delayed-match-to-sample (DMTS) task to determine how ongoing changes in brain activity corresponded with WM performance. Separate analyses were conducted for *match* trials (where target and probe orientations were the same) and *non-match* trials (where target and probe orientations were different). In match trials, participants were more likely to incorrectly report that orientations did not match when their activity patterns drifted away from the target orientation and towards target-adjacent orientations. In non-match trials, participants were more likely to incorrectly report that orientations matched when their activity patterns drifted away from the probe orientation and towards a control orientation rotated, relative to the target, in the opposite direction of the probe. These results suggest that WM failures can be at least partially explained by representational drift during maintenance. Neural drift effects analogous to those observed here have previously been theorized, and effects consistent with neural population activity drift have been observed in behavioral and modeling research; however, the present study represents, to our knowledge, the first human neuroimaging study to directly demonstrate the consequences of representational drift in brain activity patterns for WM performance.

### Consequences of representational drift in match and non-match trials

Representational drift for match trials assessed whether participants’ WM representations drifted towards the target orientation or target-adjacent orientations. Drift for accurate and inaccurate trials was similar at encoding but then quickly diverged; activity patterns drifted closer to target-adjacent orientations for inaccurate trials than accurate, suggesting that participants were more likely to incorrectly report that orientations did not match when their WM representations were more similar to target-adjacent orientations. Interestingly, representational drift showed a quadratic trend, with maximal differentiation between correct and incorrect trials in the early-to-mid maintenance period; drift indices were not significantly different between accurate and inaccurate trials at encoding or probe. (Note that effects were typically maximal at the middle of the time period we analyzed, but the analysis period terminated at probe presentation; thus, after accounting for BOLD lag, the center of the analysis period represented brain activity corresponding to ∼4s into the 11s maintenance period.) This suggests that even with successful encoding, disruption of WM patterns during the maintenance period could lead to an incorrect response on the subsequent probe. Furthermore, it did not appear necessary for this disruption to persist into the probe period to have an effect on behavior.

Representational drift for non-match trials measured whether participants’ ongoing WM representation was relatively more similar to the probe orientation or a control orientation also adjacent to the target, but in the opposite direction from the probe. Generally, representational drift for accurate and inaccurate trials was similar throughout the trial except for the maintenance period, where representations drifted towards the probe for inaccurate trials but towards the control orientation for accurate trials. This suggests that when participants’ WM representation of the target was more similar to the probe orientation, they were more likely to incorrectly report that probe and target orientations matched. As in match trials, representational drift diverged between accurate and inaccurate trials primarily during the early-to-mid maintenance period; there was no significant difference in representational drift by accuracy at either encoding or probe.

Both these results suggest that the early-to-mid maintenance period is critical to WM accuracy (in line with long-term memory findings; Ranganath, Cohen and Brozinsky, 2005; Bergmann et al., 2013) and pattern similarity at encoding and probe may not necessarily guarantee accuracy if WM patterns are disrupted during maintenance. Our findings may indicate that participants’ WM representations are most labile in the first few seconds following encoding; although in the present study, representational drift seemed to stabilize later in the maintenance period, early drift activity still had an effect on WM accuracy. Past research has documented a form of “activity-silent” WM, wherein neural activity for an unattended item drops to baseline during the maintenance period, even when the item is later successfully remembered at probe (Stokes 2015; Rose et al. 2016; Sprague et al. 2016). It is possible, then, that participants in our study actively maintained representations in early-to-mid maintenance, and then allowed those representations to become dormant, which could account for the lack of differentiation in WM drift between accurate and inaccurate trials at probe. However, it appears that even if participants’ representations became relatively activity-silent by probe time, their behavioral decisions may have been based on their WM representations from earlier in the maintenance period. In turn, this suggests that those first few seconds of WM maintenance may comprise a critical consolidation period, after which the representations become more crystallized. This reflects findings in long-term memory research, where memories for events are labile during only a limited period when the memory is active (Nader et al. 2000; Lee 2009).

### Accuracy-based differences in “raw” pattern similarity to various task-relevant orientations

While our primary hypotheses required calculating the novel representational drift indices described above in order to capture effects associated with specific WM representations (e.g., drift towards the probe orientation), we also observed differences by accuracy in the raw pattern similarities (a measure more commonly used in previous studies) used to compute those indices. In match trials, raw pattern similarity values between ongoing brain activity and any task-relevant orientation (i.e., the target orientation for that trial and the two target-adjacent orientations used as controls; see **Figures 2A–B**) were generally greater for inaccurate trials; while this varied over the timecourse of the trial (for instance, the difference was statistically significant at probe, but not at encoding or maintenance), pattern similarity was numerically greater at every timepoint throughout the trial. In other words, when pattern similarity was greater for any orientation, participants tended to report a non-match. Conversely, in non-match trials, raw pattern similarity values to any task-relevant orientation (i.e., the target orientation, the non-matching probe, and the probe-opposite control orientation; see **Figures 3A–C**) were generally greater for accurate trials. Again, this varied over the timecourse of the trial, but pattern similarity was numerically greater at most timepoints throughout the trial; this means that, as with match trials, when pattern similarity was greater for any orientation, participants tended to report a non-match. One likely explanation for this pattern of results is that higher pattern similarity to task-relevant orientations, in a manner that is not particularly specific to any one orientation or portion of the trial, reflects general alertness or task-focused states of mind; in other words, a participant who is staying focused on performing the WM task is likely to present brain activity patterns more similar to *any* task-relevant orientation than a participant who is distracted or otherwise inattentive.

If this is the case, then our pattern of results suggests an overall task strategy wherein participants tend to report non-matches more often when their attention is more focused and their representations of the WM target are clearer. Conversely, when participants’ attention was less focused and thus they had representations of the target that were less clear, they may have been more likely to report a match. Put another way, it seems probable that participants were adopting a violation-detection strategy in which they tended to report a non-match when their WM representations were sufficiently clear to establish confidence in their decision, whereas a match response could occur either because they were actually confident of a match, or because their WM representations were not clear enough to be confident of detecting a non-match. Thus, these findings based on raw pattern similarities may offer some insights into participants’ strategies for performing the WM task, although they were not particularly effective for isolating representation-specific effects and instead appeared primarily to reflect overall attention or task focus. Rather, for representation-specific effects, the *difference* between pattern similarity timelines provided the critical measure, namely, the representational drift index described in the section above.

## Conclusion

What are the causes of WM failure? In sum, our results constitute neural evidence that representational drift is among the factors that underlie such failure. When an item is held in WM, its representation is subject to random fluctuations. If those fluctuations bring the representation closer to those of non-target items that may also appear in the environment, WM errors can occur.

## Acknowledgements

We thank Lauren Bandel, Aaron Halvorsen, Rafay Khan, Joanne Murray, and Kerry Hartz for assistance with data collection and scanning.

